# Confounding factors in assessing the enriched expression of somatic mutant allele in bulk tumor samples

**DOI:** 10.1101/2025.05.27.656366

**Authors:** Kohei Hagiwara, Andrew Thrasher, Nadezhda V. Terekhanova, Jinghui Zhang

## Abstract

Allele-specific expression (ASE) of somatic mutations can be caused by *cis*-activation of the mutant allele or silencing of the wildtype allele and has been investigated by examining the enrichment of mutant allele in RNA relative to DNA. Here we show that this mutation-based approach can be confounded by gene expression differences in tumor and normal cells which co-exist in most bulk tumor samples. We modeled mutant allele expression by incorporating tumor/normal expression difference, mutant allele dosage, tumor purity, and nonsense-mediated decay (NMD) efficiency, projecting that the enrichment can occur without ASE. This confounding effect is exacerbated with low tumor purity and is dependent on mutant allele dosage for NMD-sensitive mutations. The model was validated empirically using somatic insertions/deletions (indels) from 9,101 the Cancer Genome Atlas (TCGA) samples, showing three-fold higher enrichment in driver genes vs. non-drivers. To explore if the enriched mutant allele expression can be leveraged to complement DNA-based somatic mutation detection, we performed *de novo* somatic indel calling using TCGA RNA-Seq, increasing the TCGA driver indel repertoire by ∼14%, mostly in samples with lower tumor purity. Our study not only revealed confounding factors in ASE analysis but also demonstrated the utility of leveraging RNA-Seq for driver mutation analysis.

## 1. Introduction

In a diploid genome, allele specific expression (ASE) refers to a phenomenon of preferential expression of one allele over the other in a cell, which can unveil *cis*-acting regulatory mechanisms of attenuating (including silencing) or enhancing (including activation) the transcription of one specific allele [1]. A conventional approach to characterize ASE is to test if the non-reference allele fraction (also known as the variant allele fraction, VAF) in RNA (VAF^RNA^) is significantly deviated from its expected VAF in DNA (VAF^DNA^). In non-cancerous tissues, the analysis of allelic imbalance of germline heterozygous single-nucleotide polymorphisms (SNPs) in RNA, which measures a significant deviation from the expected VAF^DNA^ of 0.5, has led to the identification of genes or regions that exhibit ASE [2]. However, when applying the same technique to cancerous tissues [3–5], these imbalances were mostly (84.3%) attributed to somatically acquired copy-number alterations (CNA) in the tumor DNA [3]. More recent studies have attempted to characterize ASE events independent of somatic CNA by focusing on ASE of individual genes (gene-based)[6] or somatic mutations (mutation-based) [7–9]. The gene-based approaches determine ASE by analyzing heterozygous germline SNPs in the transcriptome but can be constrained by marker availability. The mutation-based approaches directly compare a mutation’s VAF^RNA^ or comparable metrics to that in the tumor DNA genome and a significant deviation is attributed either to ASE or, in the case of truncating mutations, to nonsense-mediated decay (NMD). Such interpretation is valid under the condition that gene expression level is comparable in tumor cells and “contaminating” normal cells that co-exist in bulk tumor samples. However, it is currently unclear how the difference in expression levels between tumor and normal cells affects the assessment of somatic mutant allele expression relative to its VAF^DNA^. Tumor and normal tissues are known to exhibit differential gene expressions [10]. For example, the tumor suppressor gene *CDKN2A* was noted to have significantly high expression in tumor tissues compared with the matching normal tissues acquired from cancer patients [11]. Furthermore, tumor cells can undergo hypertranscription, a global change on transcription activity [12–14], which can also result in expression difference.

Here, we present a theoretical framework to investigate the impact of expression difference on the somatic VAF imbalance in bulk RNA-Seq samples. We modeled the ratio of VAF^RNA^ to VAF^DNA^, termed allelic expression variation (AEV) for convenience, with tumor purity, somatic CNA, expression difference in tumor and normal tissues, and NMD efficiency as parameters (*Modelling* workflow in **Fig. 1**). Importantly, the model does not assume ASE, i.e. an equal expression of the mutant and the wildtype alleles. Theoretical and simulation analyses of this model showed that unbalanced AEV (i.e., AEV≠ 1) can occur due to expression differences in tumor and normal cells, and that this confounding effect is more pronounced with lower tumor purity. Furthermore, for truncating mutations that can cause NMD, the mutant allele dosage can affect AEV. By focusing on mutations with enriched allelic expression (i.e., AEV > 1), we validated this theoretical model using 9,101 The Cancer Genome Atlas (TCGA) [15] samples with whole-exome sequencing (WES) and RNA-Seq datasets available on Cancer Genomics Cloud (CGC) [16] (*Validation* workflow in **Fig. 1**). In this analysis, we used somatic insertions/deletions (indels) identified in WES by the Genomic Data Commons (GDC) [17] multi-caller pipeline to enable assessing the effect of NMD and to compare AEV in tumor suppressor genes and oncogenes with that of non-driver genes. While the tumor/normal expression difference can confound the detection of true ASE, elevated AEV can be exploited for RNA-based somatic mutation detection to complement the DNA-based detection. This may be particularly effective for somatic indel detection due to a limited sensitivity by DNA-Seq at 40–50% [18, 19] (*Application* workflow in **Fig. 1**). Indeed, our *de novo* indel detection using TCGA RNA-seq data has yielded an additional ∼14% driver indels compared to the DNA-only based approach by GDC. Notably, one third of the oncogenic in-frame indels detected exclusively by this approach were rated actionable by COSMIC Actionability database [20], highlighting the potential for clinical impact.

**Figure 1.**
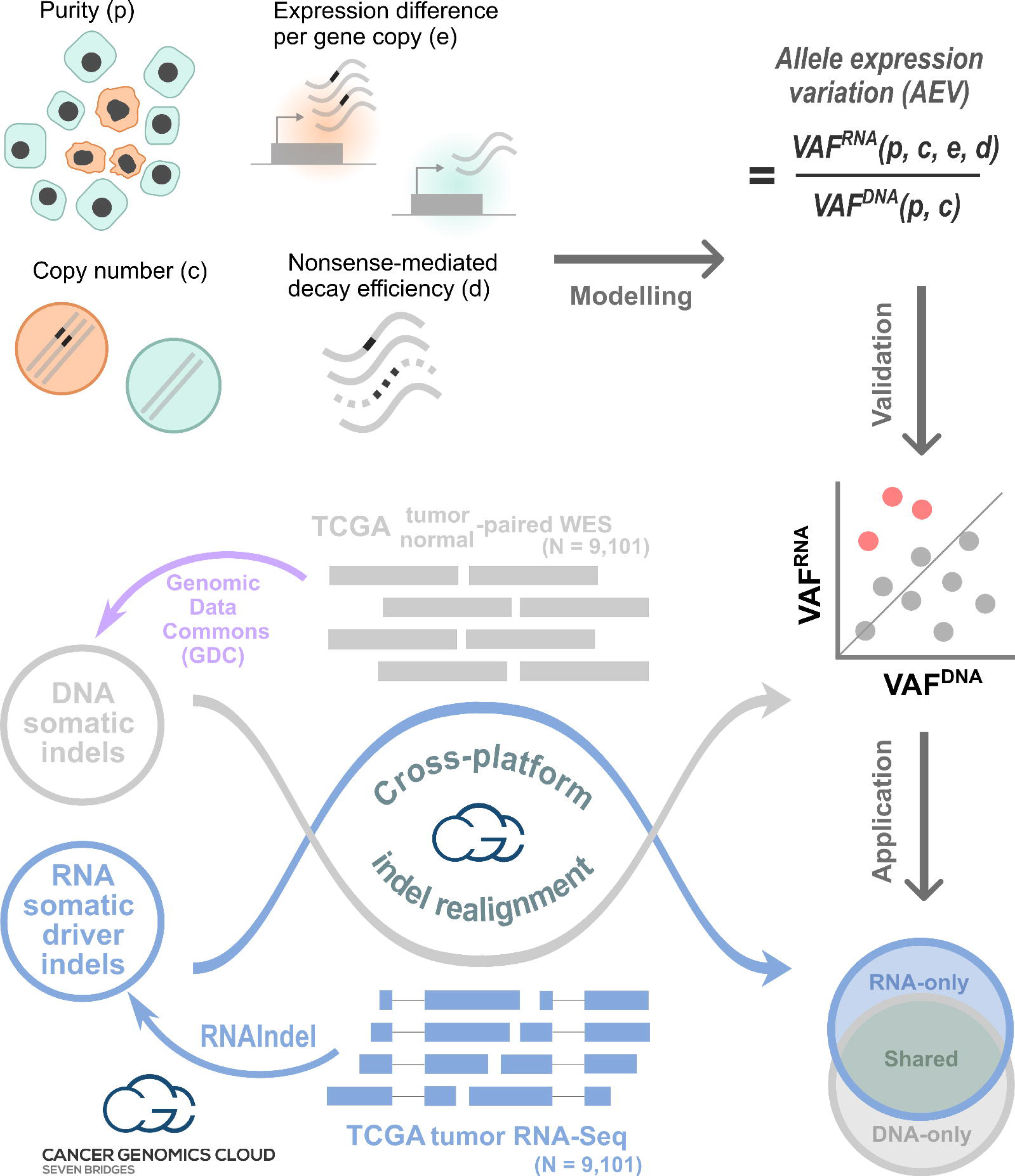
Study design with three main workflows. ***Modelling***: purity, copy number, expression level per copy in tumor relative to normal, and nonsense-mediated decay efficiency were used to model the VAF ratio of RNA to DNA, which is termed allelic expression variation (AEV) for convenience. Theoretical and simulation study was conducted to identify conditions for AEV > 1 (upward AEV). ***Validation***: DNA somatic indels identified by Genomic Data Commons’s (GDC) multi-caller platform in the TCGA dataset (purple workflow) were used to validate the model for upward AEV (red dots). Reads supporting indels were realigned in both WES and RNA-Seq to accurately calculate AEV. ***Application***: Confounding effect led to elevated VAF^RNA^ relative to VAF^DNA^ can be exploited to enhance mutation calling from RNA-Seq. This was tested for somatic driver indel detection by calling from RNA-Seq with WES read support to complement the TCGA indel set (Venn diagram). Indels in each Venn section were examined for AEV and purity along with potential clinical implications. Major computation analyses in ***Validation*** and ***Application*** were performed on Cancer Genomics Cloud.

## 2. Materials and Methods

### TCGA dataset

This study used the TCGA dataset available as GDC v36 on Cancer Genomics Cloud (CGC) (https://www.cancergenomicscloud.org). We included TCGA cases for which all of the following files were available: tumor/normal-paired WES and tumor RNA-Seq BAM files and mutation annotation format (MAF) files (n = 9,101). The MAF files were prepared by GDC by applying MuTect2 [21], VarScan2 [22], and Pindel [23] to the paired WES for somatic indel calls. Gene expression data by STAR Count [24] and copy-number data by ASCAT [25] were also available on CGC. The TCGA tumor purity data was obtained from TCGA PanCanAtlas (https://gdc.cancer.gov/about-data/publications/pancanatlas).

### Derivation of VAF ratio between RNA and DNA

In a bulk tumor sample consisting of *x* tumor and *y* normal cells, we denote *q* as:

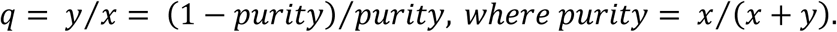

For an autosomal gene harboring a somatic mutation in tumor cells, we expect *m* and *w* copies of mutant and wildtype alleles, respectively. *m* is a positive integer and *w* is a non-negative integer, where *w* = 0 represents loss of the wildtype allele due to loss-of-heterozygosity (LOH). The normal cells are diploid and homozygous for the reference allele. The DNA VAF in the bulk tumor sample is given by:

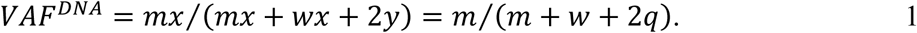

Consider that each gene copy in tumor cells is expressed at *e*-fold relative to normal, regardless of the mutation status of the allele (i.e., no ASE). The mutant allele triggers NMD with decay efficiency *d* ∈ [0, 1], where *d* = 0 represents a complete evasion from NMD and *d* = 1 represents a complete decay of the transcript. The RNA VAF is:

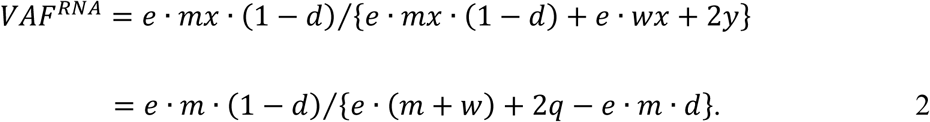

The VAF ratio of RNA to DNA (i.e., somatic mutant allelic imbalance) is:

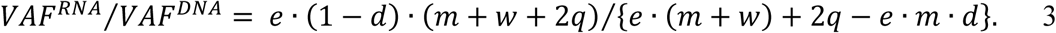

When the VAF ratio is greater than one (i.e., upward imbalance), solving for *e*, we have:

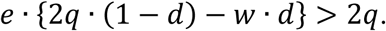

Because *e* and *q* are positive, it follows that 2*q* · (1 − *d*) − *w* · *d* > 0. As *w* and *d* are non-negative, *e* must stratify the following for upward imbalance:

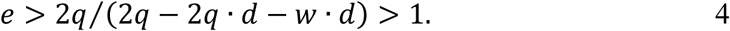

In the absence of NMD (i.e., *d* = 0), Eq.3 becomes:

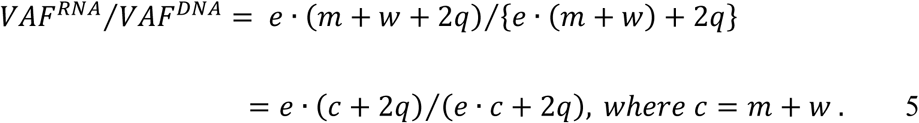

Thus, the ratio depends on the copy number *c*, but not the genotype. For example, when the copy number in the tumor cells is three (i.e., *c* = 3), AEV is identical in all genotypes: *m*: *w* = 1: 2, 2: 1, or 3: 0. In the presence of NMD, solving Eq.4 for *d*, we have the following as the efficiency upper bound for upward AEV:

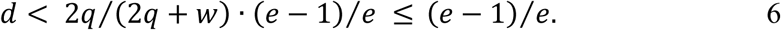

### Indel allele count benchmarking

The Sequencing Quality Control Phase 2 (SEQC2) Consortium established a reference somatic call set from the HCC1395 breast cancer cell line and the HCC1395BL cell line, which was derived from the normal B-cell of the same donor [26]. We downloaded the reference indel call set (https://ftp-trace.ncbi.nlm.nih.gov/ReferenceSamples/seqc/Somatic_Mutation_WG/release/v1.2.1/). The consortium also generated whole-genome sequencing (WGS) data on a series of diluted samples by mixing *i* parts of DNA material from HCC1395 and *j* parts from HCC1395BL for *i*: *j* = 1: 0 (undiluted), 3: 1, 1: 1, 1: 4, 1: 9, and 1: 19 [27] (https://ftp-trace.ncbi.nlm.nih.gov/ReferenceSamples/seqc/Somatic_Mutation_WG/data/SPP/). The true VAF of somatic mutations is unknown. Therefore, we used the dilution ratio as ground truth value. Specifically, the following equation holds for the amount of mutated DNA material between a sample at *n*: *m* and the undiluted sample:

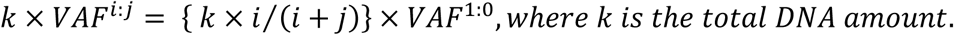

With this relation, the dilution ratio ideally matches the observed VAF ratio:

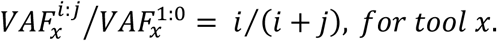

Using the serial dilution WGS dataset, we compared the variant callers used in the GDC pipeline [17] against indelPost (v0.2.3) [28] for the dilution ratio estimation. For VarScan2 (v2.3.9) [22], we used the *pileup2indel* subcommand with *--min-coverage 1 --min-reads2 1* and *- -min-var-freq 0.0001* to increase sensitivity. Pindel (v0.2.4) [23] was used with *-insert-size 400* as estimated [27]. MuTect2 (v4.1.8.0) [21] was used with the default settings. indelPost’s *count_alleles()* function was applied with the *by_fragment* option, which merges forward and reverse reads if they are from the same fragment. At 1:0, indelPost detected all the high confidence indels (n=1,467), while 1,445 were found by MuTect2, 1,196 by VarScan2, and 1,432 by Pindel. We used the unfiltered outputs from these callers (MuTect2, VarScan2, and Pindel) to maintain their sensitivity. Compared to these tools, indelPost consistently generated the best match to the expected ratios of 3/4, 1/2, 1/5, 1/10 and 1/20 over the dilution series (**Fig. S1A** and **Table S1**).

In RNA-Seq, the VAF from the diluted samples generally does not match the dilution ratio due to gene expression differences between the tumor and normal cells. Therefore, we used an RNA-Seq data for the Genome in a Bottle (GIAB) reference germline indels in the NA12878 lymphoblastoid cell line [29] (https://www.ncbi.nlm.nih.gov/bioproject/PRJNA389940) to evaluate the consistency of indel VAF in RNA-Seq with their genotype in DNA. We first selected expressed exonic indels covered with ≥ 1 RNA-Seq reads in the autosomal chromosomes. To avoid bias from NMD, we retained indels annotated as in-frame or in untranslated regions (UTR) if there were no co-occurring nonsense SNPs, frameshifts, or variants affecting splicing in the target gene. To avoid bias from ASE, we excluded genes identified to have ASE based on a prior ASE study which included NA12878 [30]. Consequently, the indels retained for this analysis are expected to exhibit balanced bi-allelic expressions with VAF^RNA^ values of 0.5 and 1.0 for heterozygous and homozygous genotype, respectively. The indel allele count tools were run with the same parameter settings as in the DNA benchmarking. The VAF^RNA^ estimates by indelPost were the closest to the expected heterozygous and homozygous genotypes of NA12878, which were determined by a pedigree analysis by the GIAB consortium (**Fig. S1B** and **Table S2**).

### Indel annotation

Indels were annotated for amino acid changes by VEP (v100) [31]. Those annotated as frameshift or in-frame insertion of a stop codon were collectively defined as truncation indels, while the other in-frame insertions/deletions were defined as in-frame indels. Bailey *et al.* [32] classified genes as drivers if their mutation frequencies were significant in the entire TCGA dataset (i.e. pan-cancer drivers) or in individual TCGA tumor types (i.e. cancer-specific drivers). Using this resource, we annotated genes as drivers if the model predicted the gene to be a pan-cancer driver, or a cancer-specific driver in the sample’s tumor type. Microsatellite instability (MSI) can induce mutations in driver genes regardless of their functional consequences, leading to over-annotation of driver status. To mitigate this confounding factor, we only used cancer-specific driver genes in MSI-enriched tumor types (COAD, ESCA, STAD and UCEC). Indels were annotated as drivers if they occurred in driver genes. Bailey *et al.* also provides tumor suppressor/oncogene classification in two confidence levels: “*tsg*” or “*possible tsg*”, and “*oncogene*” or “*possible oncogene*”. In the current study, these levels were combined to annotate driver indels either of tumor suppressor genes or oncogenes.

### Allele expression variation (AEV) analysis

To ensure accurate VAF estimation in RNA-Seq and WES, read counts for somatic indels in the TCGA dataset (*DNA somatic indels* in **Fig. 1**) were analyzed by a custom script run on CGC (*cross_platform_checker.py*), in which indelPost [28] was used for realignment. The mutant and wildtype allele counts in tumor WES and RNA-Seq were compared by Fisher’s exact test and were considered significantly different if the *p*-value was < 0.01 after Benjamini–Hochberg correction [33]. The effect size threshold was set at 20%, which required VAF^RNA^/VAF^DNA^ to be > 1.2 for upward AEV.

### Loss-of-heterozygosity (LOH) analysis

To characterize indels with LOH, we used the allele-specific copy number estimate by ASCAT [25] downloaded from CGC (see *TCGA dataset*). For genomic intervals, the copy number was separately estimated as major and minor alleles, where “*major_copy_num*” ≥ “*minor_copy_num*”. The TCGA indels were annotated with the ASCAT results based on the genomic interval overlap. Those located in regions with *minor_copy_num__* = 0 were considered candidates for LOH because one of the diploid alleles was lost. They were further assessed by considering the tumor purity estimates downloaded from the GDC portal (*TCGA dataset*). To evaluate the clonality of candidate indel and LOH, we adjusted VAF^DNA^ assuming that *x* tumor cells have a copy number *c* at the locus of interest and *y* normal cells are diploid. The read-coverage (*Cov*.) ratio between tumor and bulk sample is:

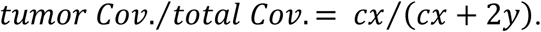

Solving for *tumor Cov*., we have:

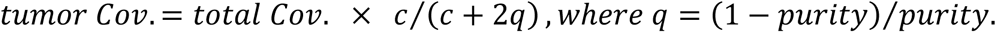

Assuming that all alternative reads are from tumor, the raw VAF^DNA^ is adjusted as:

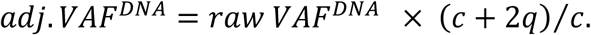

Candidate indels were inferred as clonal with LOH if their purity-adjusted VAFs were > 0.9.

### Nonsense-mediated decay analysis

We used an NMD evasion rule proposed by Lindeboom *et al.* [34, 35]. By this rule, an mRNA transcript containing premature stop codon (PTC) is classified as NMD evasive if the PTC position is <150-nt downstream of the start codon, in an exon spanning >407-nt, <55-nt upstream of the last exon junction, or within the last exon, otherwise the PTC is considered sensitive to NMD. For each frameshift, we searched for the first downstream occurrence of premature termination codon along RefSeq isoforms [36] using a custom script (*premature_stop_codon_locator.py*) and annotated the frameshifts for the NMD features based on the PTC position.

### Relative expression difference estimation

The tumor versus normal expression ratio per gene copy (variable *e* in the AEV model) was estimated under two scenarios: differential gene expression and hypertranscription. For differential expression, we first divided the gene’s transcript per million (TPM) in the tumor sample by its copy number as estimated by ASCAT [25]. As autosomal chromosomes in normal cells are diploid, we halved the median TPM from tissue-matched normal expression data also taken from TCGA for technical consistency. We used the copy-number adjusted ratio of TPM in tumor to normal median as an estimate for differential expression. Cancer types with <5 normal matching RNA-Seq data were excluded in this analysis. Zatzman *et al.* [12] estimated the degree of hypertranscription at sample level as a ratio of total RNA amount in tumor divided by that in normal with ploidy adjusted and we used their estimates for the TCGA dataset.

### Single-cell expression analysis

Single-cell or single-nucleus RNA-Seq (sc/snRNA-seq data) and cell type annotation were obtained from the previous study [37]. The methods for data normalization, feature selection, dimensionality reduction and data merging were the same as used in the study. Average gene expression visualization for *CDKN2A* and *RB1* in the selected cell groups was performed using DotPlot function from the Seurat package (v5.1.0) [38].

### Somatic indel calling from RNA-Seq

*De novo* indel calling was carried out by running RNAIndel [39] on CGC. Computing cost was a key consideration on this platform, so we refactored RNAIndel by replacing the original realignment module with the much more efficient indelPost [28]. This refactored version ran ∼10 times faster (median run time: 21 vs. 208 minutes/sample) and achieved an ∼85% reduction in computing cost (median cost: 0.16 vs. 0.97 USD/sample) (**Table S3**). We deployed this code (v.3.3.0) to CGC for the full analysis, and the cost ranged from 0.04 to 14.8 dollars/sample (median: 0.21). On CGC, RNAIndel (v.3.3.0) analysis on tumor RNA-Seq BAM files used the default parameters except for the parallelism option which was set to 12. In the output VCF files, putative indels were classified as somatic, germline, or artifactual based on the machine-learning model implemented in RNAIndel. Those classified as somatic were extracted from the VCF file and annotated for driver status (*Driver annotation*). Candidate driver indels were verified based on DNA read support, which required realignment in the corresponding paired WES data using indelPost imported in the custom script (*cross_platform_checker.py*).

### Actionability annotation

We focused on in-frame indels in oncogenes using the COSMIC Actionability database (v14) [20] by querying gene name and tumor type for the fields of GENE and DISEASE in the database. If a match was found, we examined if the mutation curated in the MUTATION_REMARK field matched the input in-frame indel. If a curated mutation was not specified, we considered the in-frame indel a match. For the matched in-frame indels, we retrieved the actionability category curated in the ACTIONABILITY_RANK field: Rank1 – approved drug demonstrated efficacy at the mutation; Rank2 – phase2/3 clinical results meeting primary outcomes; Rank3 – drug in ongoing clinical trials; Rank4 – case studies only.

### Assessment of true ASE events

We adopted a simulation-based *p*-value computation developed for somatic silencing of the *BARD1* gene in neuroblastoma [40]. Briefly, using heterozygous single-nucleotide polymorphisms (SNP) annotated as common by dbSNP [41], we calculated the sum of deviation from the heterozygous VAF (i.e.,0.5) in RNA-Seq:

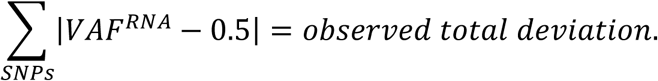

We simulated the total deviation by a binomial model with RNA-Seq coverage at each SNP locus:

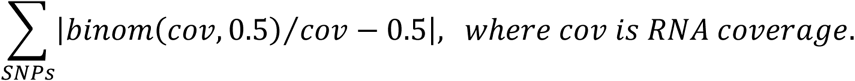

We counted the number of simulations deviating more than the observed value. The *p*-value was calculated by dividing the count by the total number of simulations (n = 1,000).

### Statistical analysis

For count data, Fisher exact and χ^2^ tests were used for pairwise and three-group comparisons, respectively. To compare distributions, Mann–Whitney and Kruskal–Wallis tests were used for pairwise and three-group comparisons. All tests were two-sided and were taken significant at *p*-value < 0.05.

## 3. Results

### Modelling allele expression variation (AEV) for somatic mutation

We considered a somatic indel with a total copy number of *c* represented by *m* and *w* copies of mutant and wildtype alleles, respectively (i.e., copy number *c* = *m* + *w*) and modeled its DNA VAF in a bulk sample (VAF^DNA^, Eq.1 in Materials and Methods). We also modeled the indel’s VAF in RNA where the gene expression per copy is *e* times different between tumor and normal, and the mutant allele is degraded at NMD efficiency *d* (VAF^RNA^, Eq.2). The ratio of VAF^RNA^ to VAF^DNA^ , defined as AEV (Eq.3) (illustrative example in **Fig. 2A**), is used to show enriched (upward AEV) or depleted (downward AEV) mutant allele representation in RNA-Seq. Our model projects that higher expression in tumor per gene copy (*e* > 1) is required for upward AEV (Eq.4) and we simulated the model over tumor purity in copy number *c* = 2 (diploid) at various NMD efficiencies (**Fig. 2B**). For mutations that do not trigger NMD (i.e., *d* = 0), no mutant allele expression variation is expected (i.e., AEV = 1) when the gene expression levels per copy are identical in tumor and normal cells (i.e., *e* = 1) in contrast to the upward and downward AEV expected for *e* > 1 and *e* < 1, respectively. Further, this effect on AEV is more prominent with lower purity due to a different rate of VAF^RNA^ reduction compared to that of VAF^DNA^ (**Fig. 2C**). In absence of NMD, the mutant genotypes have no effect on AEV (*d* = 0 in **Fig. 2B** and Eq.5). However, with *d* > 0, enriched mutant allele dosage due to CNA or LOH results in higher AEV (e.g., mut./mut. vs. mut./wt. in the three panels with *d* > 0 in **Fig. 2B**).

**Figure 2.**
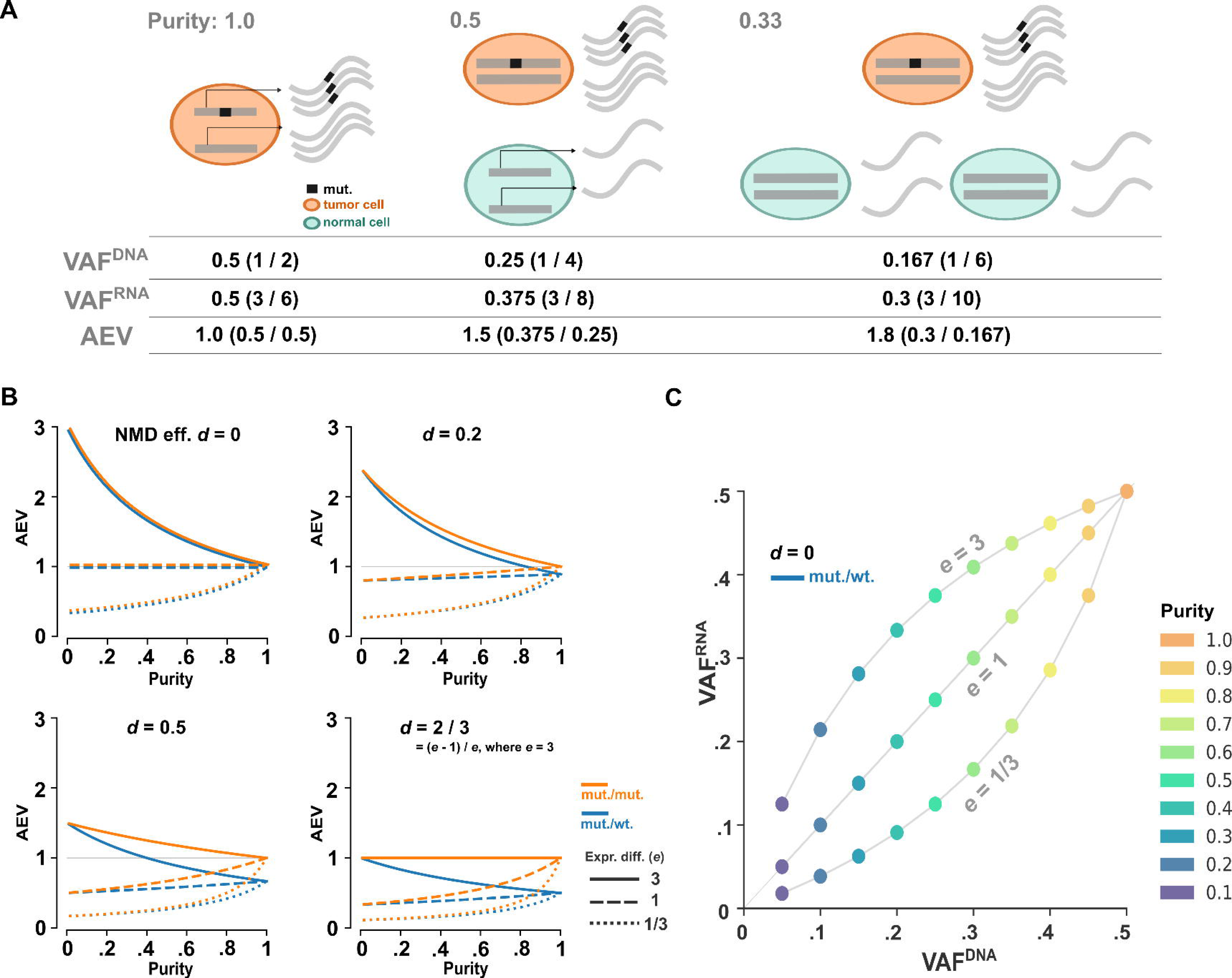
Simulation on AEV in absence of ASE. **A)** Illustration of upward AEV under the scenario that gene expression in tumor is 3-times of that in normal. In tumor cells, three copies of the mutant and wildtype alleles are transcribed while a single copy of the alleles is transcribed in normal cells. The VAF^DNA^, VAF^RNA^, and the ratio of VAF^RNA^ to VAF^DNA^ (AEV) projected from tumor purity at 1.0, 0.5 and 0.33 are presented below. **B**) Simulation of the AEV model for mono-(blue) and bi-allelic (orange) mutation in a diploid region with varying degree of tumor purity and NMD efficiencies. **C)** Simulated VAF^DNA^ vs. VAF^RNA^ under three expression difference scenarios for purity ranging from 0.1–1.0. The simulation is performed for NMD-insensitive, mono-allelic mutation.

While higher NMD efficiency decreases AEV, upward AEV is still detectable if the efficiency is smaller than (*e* − 1)⁄*e* (e.g., *d* = 2⁄3 in **Fig. 2B** and Eq.6). Similar results are also obtained in other copy number states (**Fig. S2**).

In summary, under the condition where the gene expression per copy is higher in tumor than in normal cells (i.e., *e* > 1), upward AEV (i.e., AEV > 1) can be detected in absence of ASE and the pattern is more pronounced with reduced tumor purity. While this pattern is more prominent with NMD-evasive mutations, AEV can be retained at a higher level with enriched mutant allele dosage.

### Upward AEV in somatic truncations in TCGA

To empirically validate the AEV model, we used 106,157 somatic coding indels on the autosomal chromosomes identified by the multi-caller pipeline developed by NCI Genome Data Commons (GDC). These indels were annotated for amino acid changes and the underlying genes were categorized as tumor suppressor genes (TSGs), oncogenes, or non-driver genes (Materials and Methods). Truncation indels (frameshifts and in-frame insertions of stop codons) accounted for the vast majority of all somatic indels (92.93% or 98,649/106,157); those in driver genes primarily disrupted TSGs (92.14%, 4,724/5,127). In TSGs, truncations are frequently accompanied by LOH, conforming to the two-hit hypothesis of tumorigenesis [42]. Therefore, this indel class is well suited for testing the predicted effects of LOH and NMD efficiency in addition to expression difference and tumor purity, which are applicable to other mutation types.

First, we quantified indel VAFs in both WES and RNA-Seq using mutant and wild-type read counts extracted by indelPost [28], as this tool outperformed those used by GDC in the benchmarking of indel VAF with serially diluted tumor DNA samples and expressed indels in RNA-Seq (**Fig. S1** and Materials and Methods). When NMD occurs at a high efficiency, the transcript harboring the truncation allele may be degraded and thus undetectable, but the wildtype transcript can be present in the transcriptome. To capture this scenario, we required the indel locus to be covered with ≥ 1 RNA-Seq reads regardless of mutation status, resulting in inclusion of 4,450 truncations in tumor suppressor genes and 69,840 in non-driver genes. Strikingly, upward AEV was four times as prevalent in tumor suppressor genes as in non-driver genes (7.69% vs. 1.76%, Fisher exact *p* < 2.2×10^-16^) (**Fig. 3A**).

**Figure 3.**
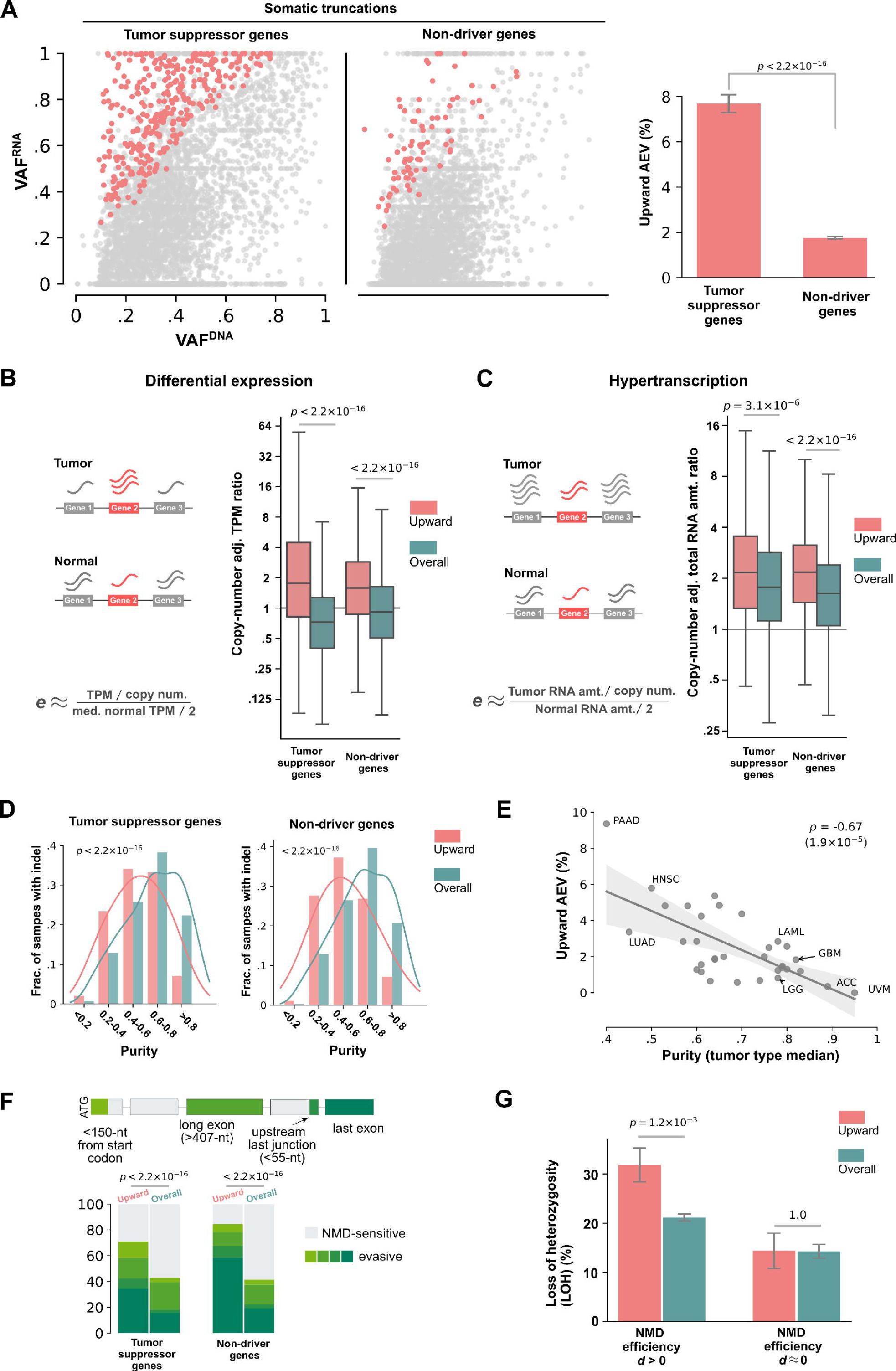
Evaluation of the AEV model using somatic truncation indels in TCGA samples. **A)** Comparison of upward AEV indels (AEV> 1.2 with FDR < 0.01) in tumor suppression genes (TSGs) and non-driver genes. *Left*, scatter plots showing VAF^DNA^ (x-axis) and VAF^RNA^ (y-axis) for indels in TSGs and non-drivers with upward AEVs highlighted in red dots. Non-driver indels were downsampled to match the TSG indel size for visualization. *Right*, comparison of upward AEV prevalence in TSGs and non-drivers by Fisher exact test. The full data set was used which contains 4,450 and 69,840 indels in TSGs and non-drivers, respectively. Evaluation of expression difference in tumor vs. normal on upward AEV is shown in **B)** for differential expression and in **C)** for global upregulation in transcription activity (hypertranscription). For measuring differential expressions in **B**), relative expression per allele (*e*) was estimated by dividing each tumor sample’s TPM with the median TPM in tissue-matched normal samples adjusted for copy number. Box plots represent the distribution of the TMP ratio for samples harboring upward AEV indels (*Upward*) with all indel-containing samples (*Overall*) with p values by Mann– Whitney test. The expression ratio plotted in **C)** was based on the RNA amount ratio estimated by Zatzman *et al.* [12]. **D)** and **E)** present the effect of tumor purity on upward AEV. **D)** Comparison of tumor purity distribution in samples harboring upward indels (*Upward*) vs all indel-containing samples (*Overall*) by Mann–Whitney test. **E)** Correlation of upward AEV prevalence (y-axis) in a tumor type with the median purity of the tumor type (x-axis) by linear regression. PAAD: Pancreatic adenocarcinoma; HNSC: Head and neck squamous cell carcinoma; LUAD: Lung adenocarcinoma; LAML: Acute myeloid leukemia; GBM: Glioblastoma multiforme; LGG; Lower grade glioma; ACC: Adrenocortical carcinoma; UVM: Uveal Melanoma. The effect of NMD on upward AEV is shown in **F)** for comparison of NMD features by χ^2^ test and in **G)** for comparison of the prevalence of LOH by Fisher exact test stratified by NMD-sensitive (*d* > 0) and evasive (d ≈ 0) indels.

We then examined whether the upward AEV cases matched the predicted pattern of model parameters: relative gene expression between tumor and normal cells, purity, copy number, and NMD efficiency. To evaluate differential expression, we estimated the *e* as the TPM ratio between tumor and tissue-matched normal controls adjusted for copy number alterations (**Fig. 3B**) (Materials and Methods). Consistent with our model, *e* was higher for genes carrying upward-AEV indels (median: 1.76 in upward vs 0.72 in overall for TSGs; 1.60 vs 0.92 for non-driver genes, Mann–Whitney *p* < 2.2×10^-16^ for both gene classes). Alternatively, the *e* variable can be > 1 under a global upregulation of transcription activity (hypertranscription), which was recently found to be widespread in human cancers [12]. Using the degree of hypertranscription in the TCGA cohort by Zatzman *et al.* [12] (Materials and Methods), we found a higher degree of transcription activity in samples harboring the upward-AEV indels (**Fig. 3C**, median: 2.17 in upward vs.1.77 in overall for TSGs at Mann–Whitney *p* = 3.1×10^-6^; 2.17 vs. 1.63 for non-driver genes at *p* < 2.2×10^-16^). Upward AEV indels tended to be found in lower purity samples (median purity: 0.53 in upward vs. 0.66 in overall for TSGs; 0.49 vs. 0.64 for non-driver genes, Mann– Whitney *p* < 2.2×10^-16^ for both gene classes), which matched the predicted trend in our model (**Fig. 3D**). Tumor purity varies greatly by cancer type; for example, tumor samples of pancreatic adenocarcinoma (PAAD) tend to have low purity due to high stromal content (50 – 80%) [43], whereas those of brain cancers often have high purity [44]. Indeed, there was a negative correlation between purity and upward-AEV prevalence across the 33 TCGA tumor types (correlation coefficient *ρ* = -0.67, *p* = 1.9×10^-5^) with the highest upward-AEV prevalence (9.38%) detected in PAAD, the cancer type with the lowest tumor purity (median purity of 0.4) (**Fig. 3E**).

We further examined conditions relevant to truncation indels. Consistent with the model, premature stop codons (PTCs) caused by upward-AEV indels occurred more frequently in NMD evasive regions (**Fig. 3F**, χ^2^ *p* < 2.2×10^-16^ for both tumor suppressor genes and non-driver genes). As expected by the two-hit hypothesis, the overall frequency of LOH was ∼10 times higher in TSGs (18.00% in TSGs vs. 1.70% in non-driver genes, Fisher exact *p* < 2.2×10^-16^) (Materials and Methods). Therefore, we used truncation indels in TSGs to examine whether higher allele dosage of NMD-sensitive mutations caused by LOH is accompanied by upward AEV as projected by our simulation (**Fig. 2A** and **S2**). We found that indeed, LOH was enriched in upward-AEV indels with PTC in NMD-sensitive regions (NMD efficiency *d* > 0 in **Fig. 3G**; 31.87% vs. 21.19%, Fisher exact *p* = 1.2×10^-3^). By contrast, LOH was not enriched when the truncation indels created PTCs in the last exon, where the NMD efficiency is close to 0 [35] (*d* ≈ 0 in **Fig. 3G**, 14.43% vs. 14.31%, *p* = 1.0). Therefore, the empirical data was consistent with the projection of our model showing an association between LOH and AEV only for d > 0 (**Fig. 2A** and **S2**).

### Upward AEV in somatic in-frame indels in TCGA

In-frame indels accounted for 7.07% of the TCGA somatic indels and are not expected to trigger NMD. To evaluate the AEV of in-frame indels in driver genes, we focused on oncogenes as in-frame indels accounted for a much higher proportion of somatic indels than in TSGs (41.51% vs. 7.86%, Fisher exact *p* < 2.2×10^-16^). Upward AEV was more frequent in in-frame indels in oncogenes than in non-driver genes (**Fig. 4A**, 20.59% vs 6.47% , Fisher exact *p* = 2.7×10^-12^) and was significantly enriched in samples with low tumor purity (**Fig. 4B**, median: 0.43 in upward vs. 0.63 in overall for oncogenes at Mann–Whitney *p* = 8.2×10^-7^). Interestingly, differential expression between tumor and normal tissues was not a significant contributor to the upward AEV in oncogenes (left in **Fig. 4C**, median: 0.88 in overall vs. 0.93 in upward at Mann–Whitney *p* = 0.51), whereas a higher degree of hyperactivation was associated with the upward pattern (right in **Fig. 4C**, median: 1.52 in overall vs. 2.48 in upward at *p* = 2.4×10^-3^). Upward AEV cases included known gain-of-function in-frame indels such as *EGFR* deletions in exon 19 and insertions in exon 20 [45], *ERBB2* duplications of Y772_A775 [46] and *CTNNB1* deletions at S45 [47]. Considering this, the lack of differential expression may suggest the possibility of bona fide ASE in upward AEV cases.

**Figure 4.**
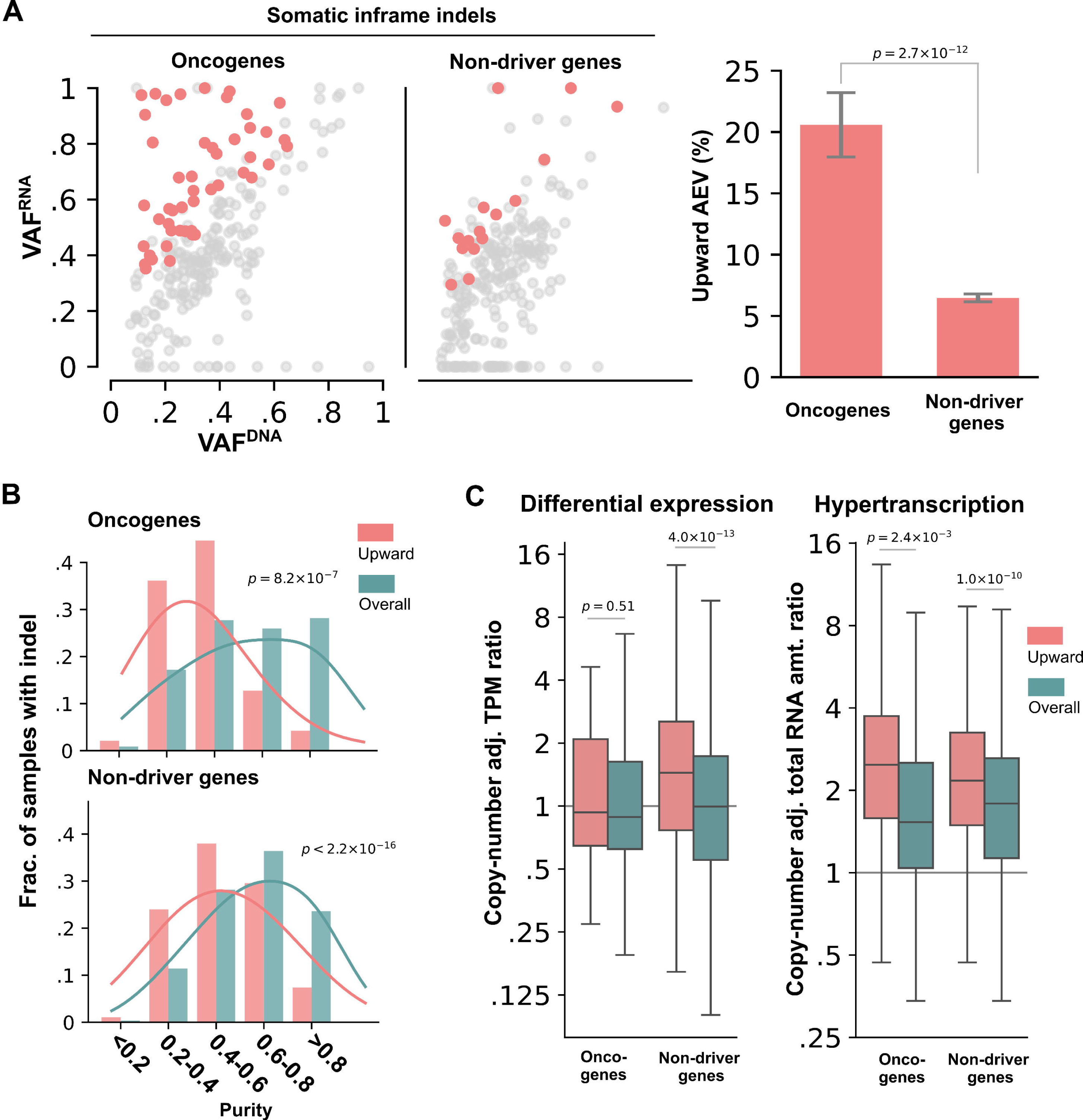
Evaluation of the AEV model using somatic in-frame indels in TCGA samples. Upward AEV prevalence, comparison of tumor purity and tumor expression ratio in upward AEV vs all indels are presented in **A)**, **B)** and **C)**, respectively, using the same style and statistical test as the corresponding analyses in Fig. 3. **A)** Comparison of upward AEV indels in oncogene and non-driver genes using down-sampled non-driver indels for a scatter plot (l*eft*) and the full data set (238 indels in oncogenes, 5,673 indels in non-drivers) for bar plot (*right*). **B)** Comparison of tumor purity distribution in samples harboring upward AEV indels (*Upward*) vs all indel-containing samples (*Overall*). **C)** Comparison of TPM ratio based on differential expression (left) and hypertranscription (right) between *Upward* and *Overall* indels.

### De novo driver indel detection in RNA-Seq

The elevated AEV in driver indels, projected by our model and validated using the TCGA dataset, suggests that RNA-Seq can complement DNA-based indel detection, particularly in samples with low tumor purity. To explore this, we performed a *de novo* indel detection in TCGA RNA-Seq focusing on truncations in tumor suppressor genes and in-frame indels in oncogenes. A total of 3,259 somatic driver indels were called from tumor RNA-Seq with ≥ 1 DNA supporting reads in the matched WES (Materials and Methods). When compared to 4,971 driver indels identified from WES data, we found that 2,390 were detected only by WES (*DNA-only* in **Fig. 5A**), 2,581 by both WES and RNA-Seq (*Shared*) and the remaining 679 detected only by RNAIndel analysis (*RNA-only*). The indels in *DNA-only* had a significantly lower VAF^RNA^ than in VAF^DNA^ (median: 0.11 in RNA vs 0.33 in DNA, Mann–Whitney *p* < 2.2×10^-16^). By contrast, the opposite pattern was found in *RNA-only* indels with much higher VAF^RNA^ (0.36 in RNA vs. 0.26 in DNA, *p* = 1.8×10^-12^) and the *Shared* indels had a slightly elevated VAF^RNA^ (median: 0.43 in RNA vs. 0.39 in DNA, *p* = 1.3×10^-11^) (**Fig. 5B**). When evaluating upward AEV, *DNA-only* had only a minimum presence (1.08%) in contrast to the significantly high fraction detected in *Shared* (13.8 %), and *RNA-only* (16.7%, χ^2^ *p* < 2.2×10^-16^) (**Fig. 5C**). Consistent with our model, *RNA-only* indels were enriched in samples with low tumor purity, for example, samples with purity < 0.4 accounted for 25.04% *RNA-only* indels in contrast to 14.69% in other categories. (**Fig. 5D**, Kruskal–Wallis *p* = 2.1×10^-8^).

**Figure 5.**
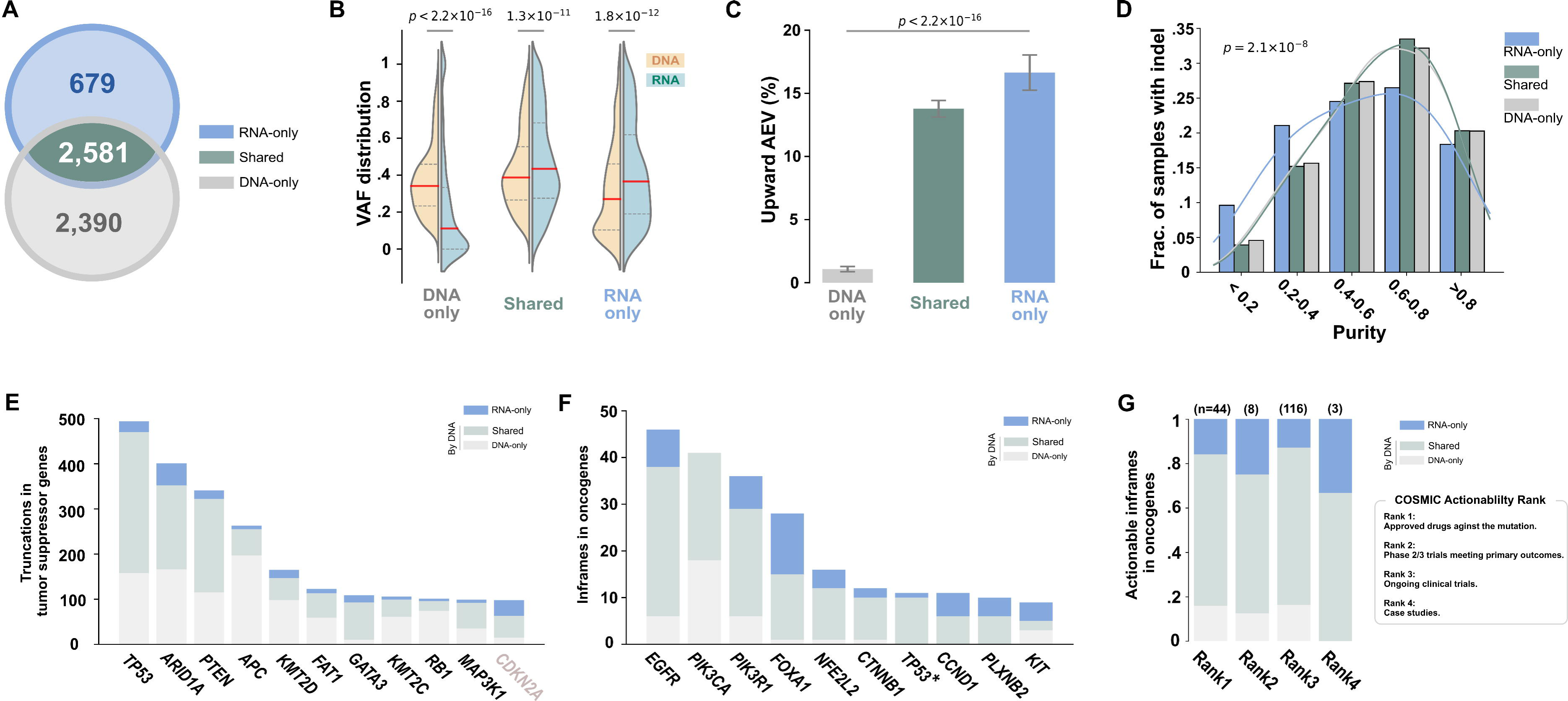
Driver indels detected by RNA-Seq and WES in TCGA samples. Driver indels are defined as in-frame mutations in oncogene or truncations in TSGs. **A)** Venn diagram showing driver indels detected exclusively by GDC’s WES analysis (*DNA-only*), by our *de novo* RNA-Seq analysis (*RNA-only*), or by both approaches (*Shared*). **B)** Comparison of VAF distributions in WES (*DNA*) and RNA-Seq (*RNA*) by Mann–Whitney *p* value for driver indels in these three categories. Median VAF is shown by the red line. **C)** Comparison of prevalence of upward AEV indels in these three categories by χ^2^ test. **D)** Tumor purity distribution for samples containing indels in the three categories. Kruskal–Wallis test was performed for assessing their differences. **E)** Top 10 most frequently mutated TSGs contributed by varying indel discovery approaches. *CDKN2A* (ranked top 11^th^) is also shown as an example for leveraging AEV for discovery due to its >20-fold higher expression in tumor (**Fig. S3**). **F)** Top 10 most frequently mutated oncogenes. *:*TP53* was annotated as “possible oncogene” in glioblastoma multiform (GBM) and low-grade glioma (LGG) by Bailey *et al.* [32]. **G)** Actionability of oncogenic in-frame indels based on COMSIC Actionability database.

The indels detected by *de novo* RNA-Seq increased truncations in TSGs by 12.74% (n = 602) compared to those known by DNA analysis (4,724). The top 10 most frequently mutated TSGs had a varying degree of contribution by RNA-Seq (**Fig. 5E**). Although not amongst the top 10, *CDKN2A* (ranked 11^th^) is a notable example of having the biggest gain of somatic indels contributed by RNA-Seq analysis, resulting in a 35.71% increase due to the 24.6-fold expression difference in tumor versus normal. By contrast, *RB1* (ranked 9^th^), which has lower expression in tumor (0.48-fold in tumor), only had a marginal gain (5.0%) by RNA-Seq. The differential expression patterns of these two genes in tumor vs. normal TCGA bulk samples were also replicated in single-cell RNA-seq data across multiple cancer types (**Fig. S3**). Notably, *de novo* RNA-Seq indel analysis reaped the biggest gain in in-frame indels in oncogenes with an overall 31.17% increase (77 added to 247 in-frame indels detected by DNA) across multiple oncogenes (**Fig. 5F**). For example, RNA-Seq indels nearly doubled driver indels in *FOXA1*, an oncogene for prostate cancer and breast cancer [32], with 13 RNA indels added to the 15 detected by WES. Of all 28 *FOXA1* indels, the vast majority (71.4%) are located between D249 and E269, which are known to be associated with favorable prognosis in prostate cancer [48]. In addition to prognostic biomarkers, 32.46% (25/77) of the additional oncogene in-frame indels were rated actionable by COSMIC Actionability [20] (**Fig. 5G**). These included targetable indels by approved drugs (Rank1 in COSMIC Actionability classification) such as *EGFR* exon 19 deletions in lung adenocarcinoma (n = 5) and *FLT3* deletion at I836 in acute myeloid leukemia (n = 2).

## 4. Discussion

In this study, we evaluated the effect of tumor/normal gene expression difference on somatic allele enrichment in bulk RNA-Seq. While a recent study was conceptually aware of such effects [8], our development of the AEV model has enabled a systematic evaluation of multiple factors contributing to the elevation of AEV. We demonstrated that elevated AEV in a bulk tumor sample can be caused by higher expression in tumor in absence of allele-specific expression.

Furthermore, this confounding effect is more pronounced in samples with low tumor purity, which, at first glance, appeared to be paradoxical (**Figs. 2A, 3D–E**). This phenomenon is due to a slower decline of VAF^RNA^ compared to that of VAF^DNA^ with lower tumor purity when gene expression is higher in tumor than normal (**Fig. 2C**). It is noteworthy that the same paradoxical trend can also arise from bona fide ASE, where the haplotype harboring the mutant allele is preferentially expressed in tumor, when examining the enrichment of mutant allele (**Fig. S4**). By contrast, if allelic imbalance of heterozygous germline SNPs were evaluated instead, the enrichment of preferentially expressed haplotype would diminish with low tumor purity as expected (**Fig. S4**).

The approach using heterozygous SNPs can also be valuable to find elevated AEV due to true ASE. For example, when tumor cells are diploid, the VAF^RNA^ of heterozygous SNPs in the bulk tumor sample remains 0.5 if the allele expression is balanced (no ASE), regardless of purity and tumor/normal expression differences. To illustrate the contrasting patterns, we show allelic imbalance of SNPs associated with *FAT1* V1689fs (AEV = 1.94) in a head-and-neck squamous cell carcinoma, which significantly deviated from 0.5 (simulation-based *p* = 0.00 in **Fig. S5**) (Materials and Methods), whereas those with *MAP3K1* P324_N325fs (AEV = 2.23) in a breast cancer did not change (*p* = 0.08). When tumor cells are not diploid, accurate measures of purity and expression difference are required as the SNP VAF^RNA^ depends on these factors. A more systematic analysis to identify upward AEV caused by ASE in the current TCGA RNA-Seq data is challenging, as the SNP-based method is only applicable to a limited number of cases having a sufficient number of heterozygous exonic SNPs in TCGA RNA-Seq data which were prepared with poly-A enrichment protocol. An RNA-seq data set generated from total RNA library preparation may potentially mitigate this limitation by leveraging the expression pattern of intronic SNPs in un-spliced pre-mRNA.

RNA-Seq has been noted to be useful for detecting low VAF mutations missed by DNA-based analysis [49]. Considering that lower purity can further amplify the relative VAF difference between DNA and RNA, the low VAF of such missed mutations might have been due to normal cell contamination rather than subclonality. Performing *de novo* mutation detection in RNA-Seq to complement DNA-based analysis would be particularly important for capturing mutations relevant for therapy. For example, in our de novo RNA-Seq indel analysis, in-frame indels in oncogene, which are often actionable, had a >30% increase compared to the DNA-based repertoire provided by GDC. Truncation indels can also be important for designing cancer immunotherapy because frameshifts may create potent neoantigens containing amino acid sequences distinct from the original [50]. Among the frameshift indels, only those evading NMD are known to be relevant to therapy [35, 51] and it is encouraging that such variants were the most prevalent in indels detected exclusively from RNA-Seq (**Fig. S6**, 41.5% in *DNA-only* vs. 49.2% in *Shared* vs. 63.1% in *RNA-only*, χ^2^ *p* < 2.2×10^-16^).

In summary, we identified tumor/normal expression difference as a confounding factor contributing to enriched somatic mutant allele expression in bulk tumor samples. The confounding effect is particularly pronounced in low-purity tumor samples. These findings serve to caution the interpretation of allelic imbalance characterized by somatic mutation alone, which suggests the inclusion of germline heterozygous SNPs for detecting bona fide ASE. At the same time, they also highlight an opportunity to exploit the use of RNA-Seq for *de novo* somatic mutation detection which can complement DNA-based mutation detection in samples with low tumor purity.

## Data Availability

This current study has not generated new datasets. Custom code developed for this study and Supplementary Tables S4–S6 has been deposited in Zenodo (https://zenodo.org/records/15492145). The refactored version of RNAIndel has been integrated into the original code repository as version 3 (https://github.com/stjude/RNAIndel).

## Acknowledgements

This study was supported by a Cancer Center Support Grant (P30 CA21765) to St. Jude Children’s Research Hospital, specifically through a Supplement (3P30CA021765-41S3) to J.Z. The study was also supported by the American Lebanese Syrian Associated Charities. We thank Mr. Michael Edmonson for his help with proofreading the manuscript.

## Supplementary Materials

**Figure S1.**
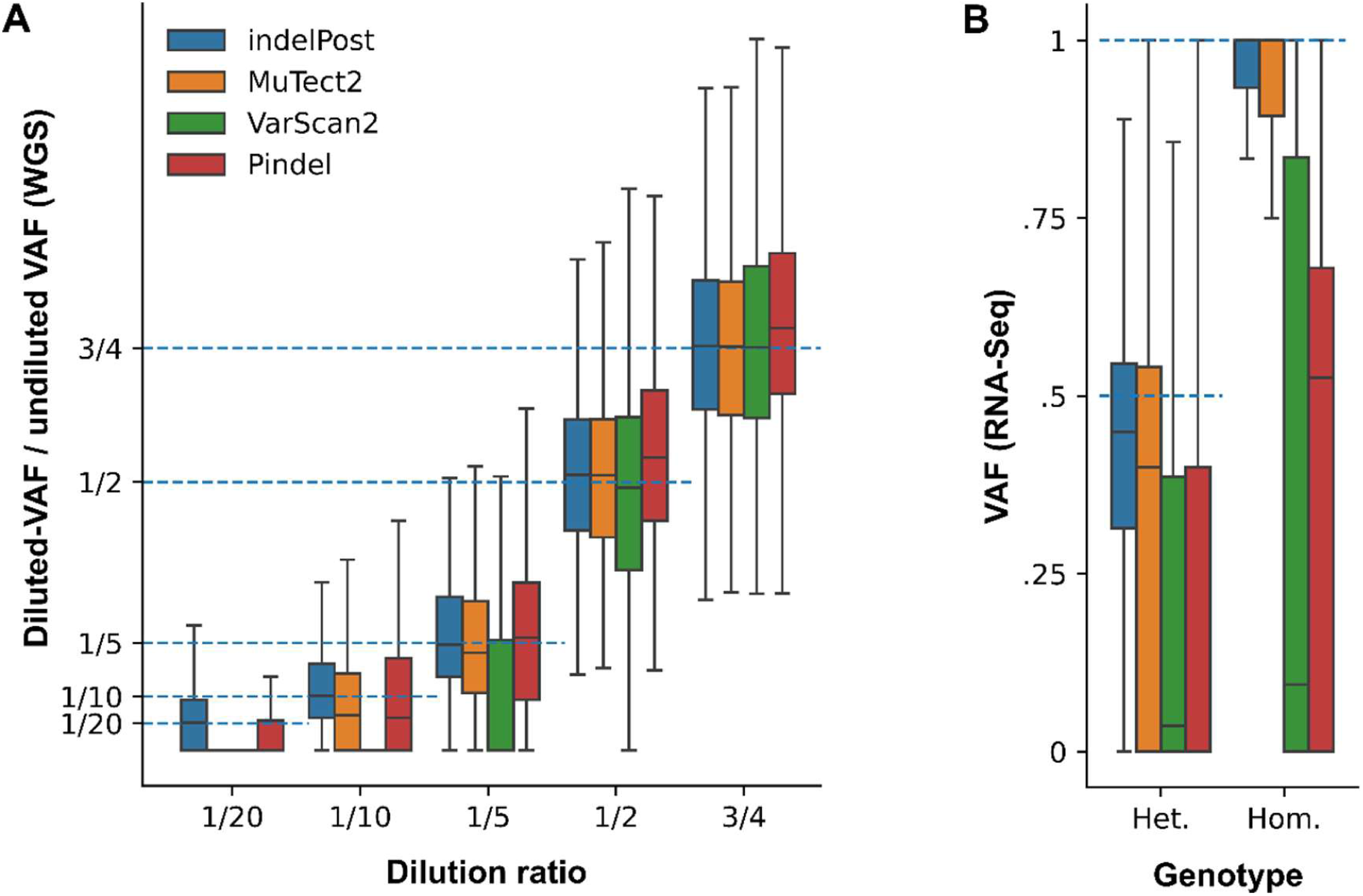
Benchmarking read counts for indel allele quantification. **A)** Quantification accuracy was compared for indelPost [1] and the indel calling tools [2–4] used by Genome Data Commons. The reference data sets include somatic indels from a tumor/normal-paired cell line (HCC1395/ HCC1395BL) with whole-genome sequencing (WGS) generated from a serial dilution experiment which has been used for benchmarking of analytical methods [5, 6]. The ground truth value was set as the ratio of VAF measured in the WGS generated from diluted samples divided by VAF in the undiluted WGS sample (see Materials and Methods for detail). Median absolute errors (MAE) are shown in **Table S1**. **B)** Reference germline indel calls from the NA12878 lymphoblastoid cell line were used for benchmark indels in RNA-Seq [7]. The ground truth was set to 0.5 for heterozygous (Het.) and 1.0 for homozygous (Hom.) genotypes as established by Genome in A Bottle. To mitigate bias from nonsense-mediated decay (NMD), indels in untranslated regions (UTRs) or in-frames were included. Also, indels in genes with known allele-specific expression (ASE) [8] in this cell line were removed from benchmarking. MAE is shown in **Table S2**.

**Figure S2.**
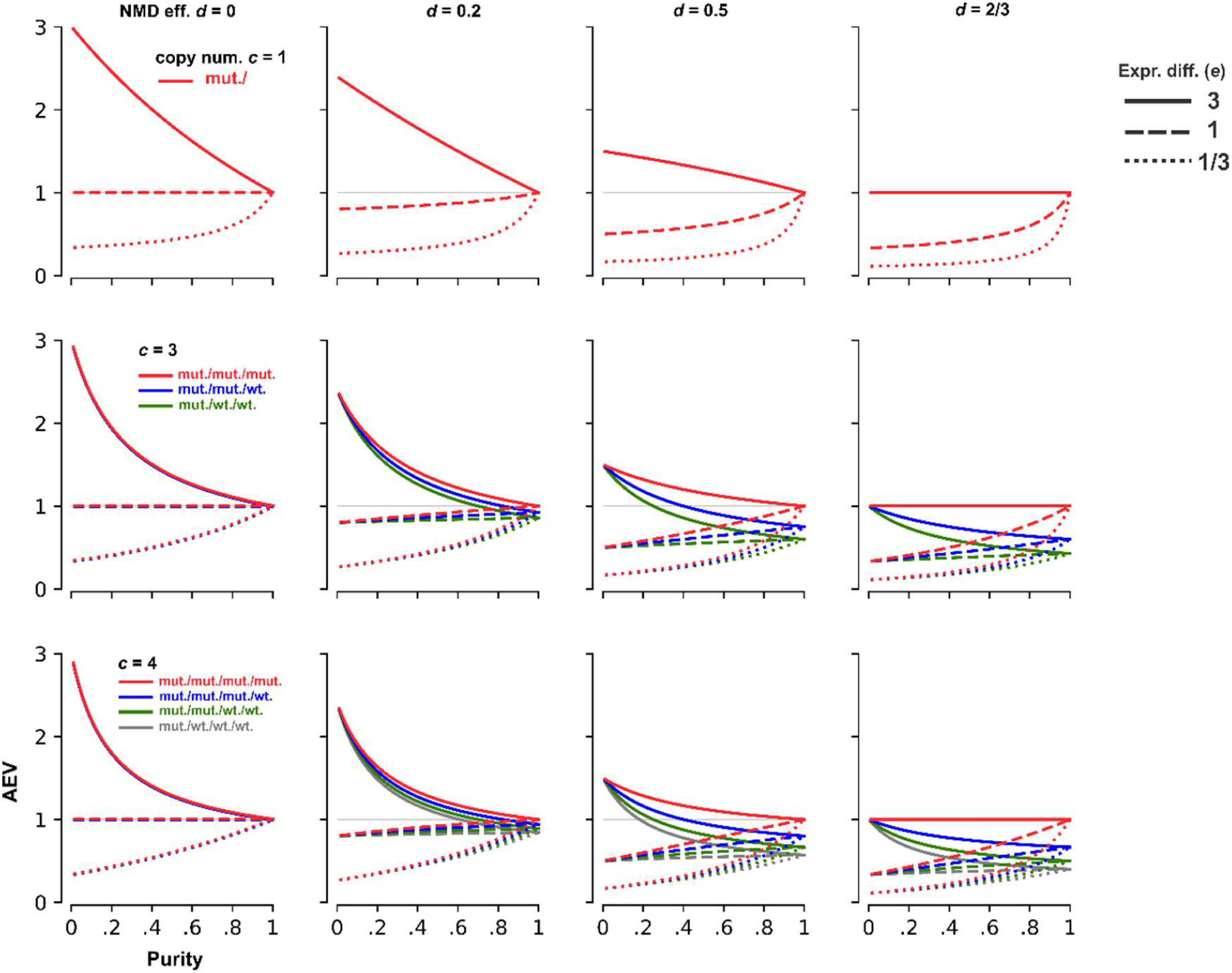
Simulation results of AEV in non-diploid regions. The AEV model is simulated for copy numbers *c* = 1 (top row), 3 (middle) and 4 (bottom) at varying NMD efficiency. Varying mutant allele dosage for each copy number category is labeled by a distinct color. The fourth column illustrates the NMD efficiency *d* = 2/3, representing the upper bound of efficiency to observe an upward AEV for copy-number adjusted expression difference *e* = 3.

**Figure S3.**
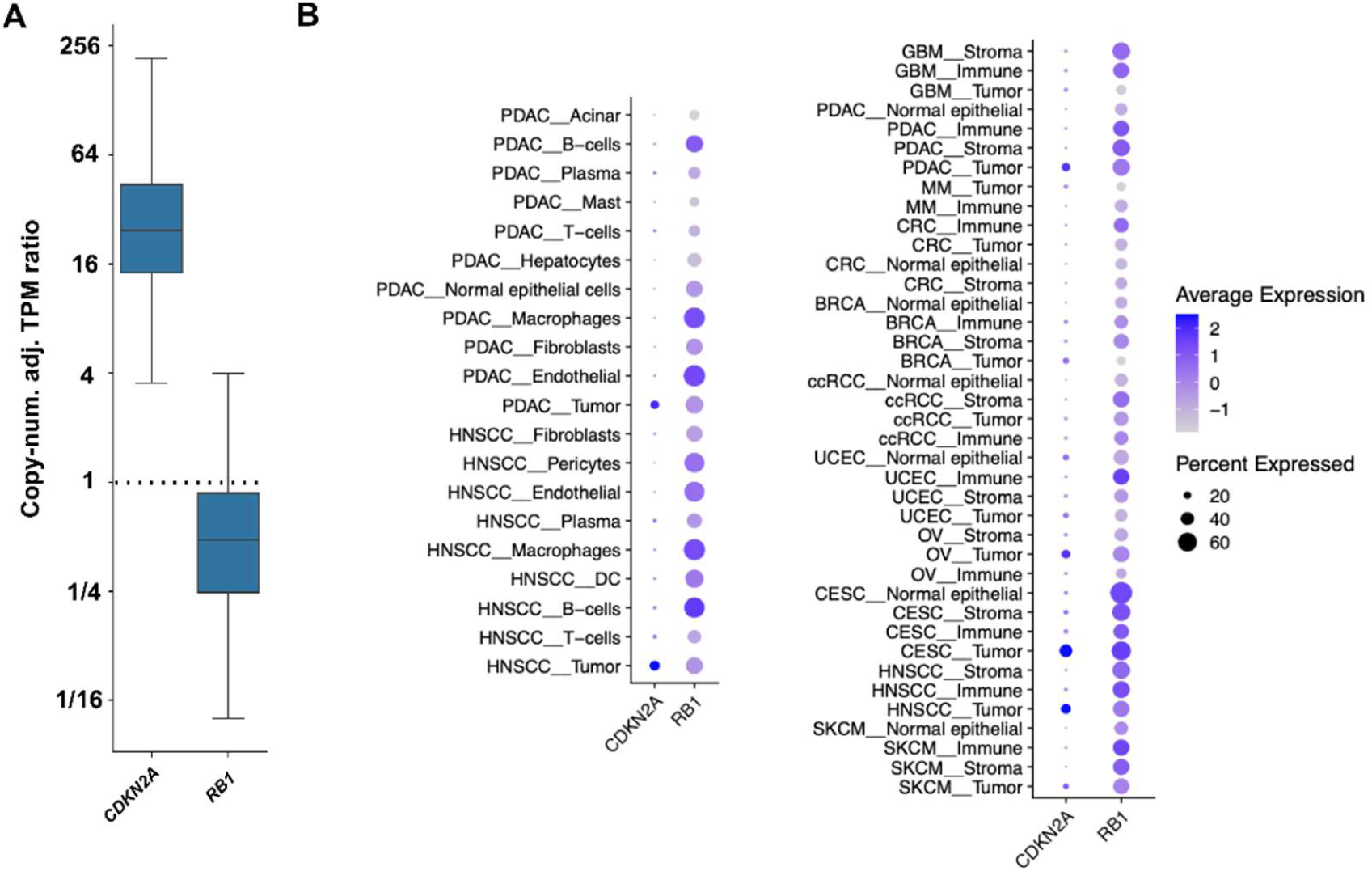
Differential expression in tumor versus normal cells for *CDKN2A* and *RB1*. **A.** The ratio of tumor vs normal expression of the bulk TCGA samples for *CDKN2A* and *RB1* (similar to main Fig. 3B). **B.** Expression of *CDKN2A* and *RB1* in tumor and normal cells based on sc/snRNA-seq from a public dataset [9]. The size and color of each circle corresponds to the percentage of positive cells and the median expression level, respectively. Left, *CDKN2A* and *RB1* expression in tumor cells and diverse types of normal cells in pancreatic ductal adenocarcinoma (PDAC) and head and neck squamous cell carcinoma (HNSCC), the two cancer types that have the lowest purity. Right, expression in tumor and normal cells across a broad spectrum of cancers. Normal cells are classified into the major categories of stroma, immune and epithelial cells (when applicable).

**Figure S4.**
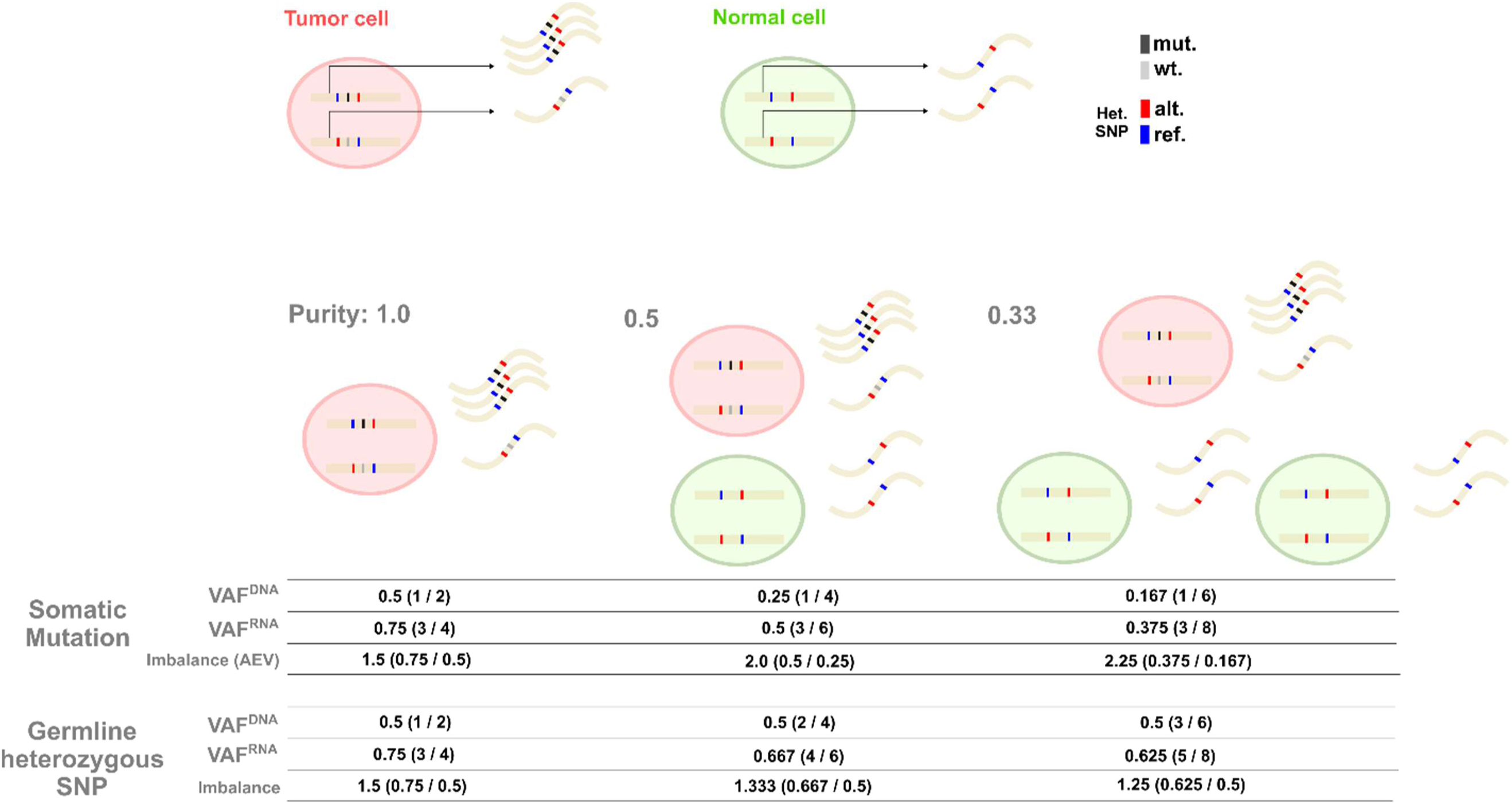
Illustration of mutation-based versus SNP-based allelic imbalance analyses. In this example, the haplotype harboring the mutant allele is transcribed 3 times as of the wild-type allele in tumor cells (true ASE) and there is no differential expression in non-mutant allele in tumor and normal cells. Allelic imbalance in this scenario is calculated in samples with purity 1.0, 0.5 and 0.33. The mutation-based approach uses the read counts of the somatic mutation to calculate the imbalance (AEV) as the ratio of VAF^RNA^ to VAF^DNA^, while the SNP-based approach uses the heterozygous SNP read counts.

**Figure S5.**
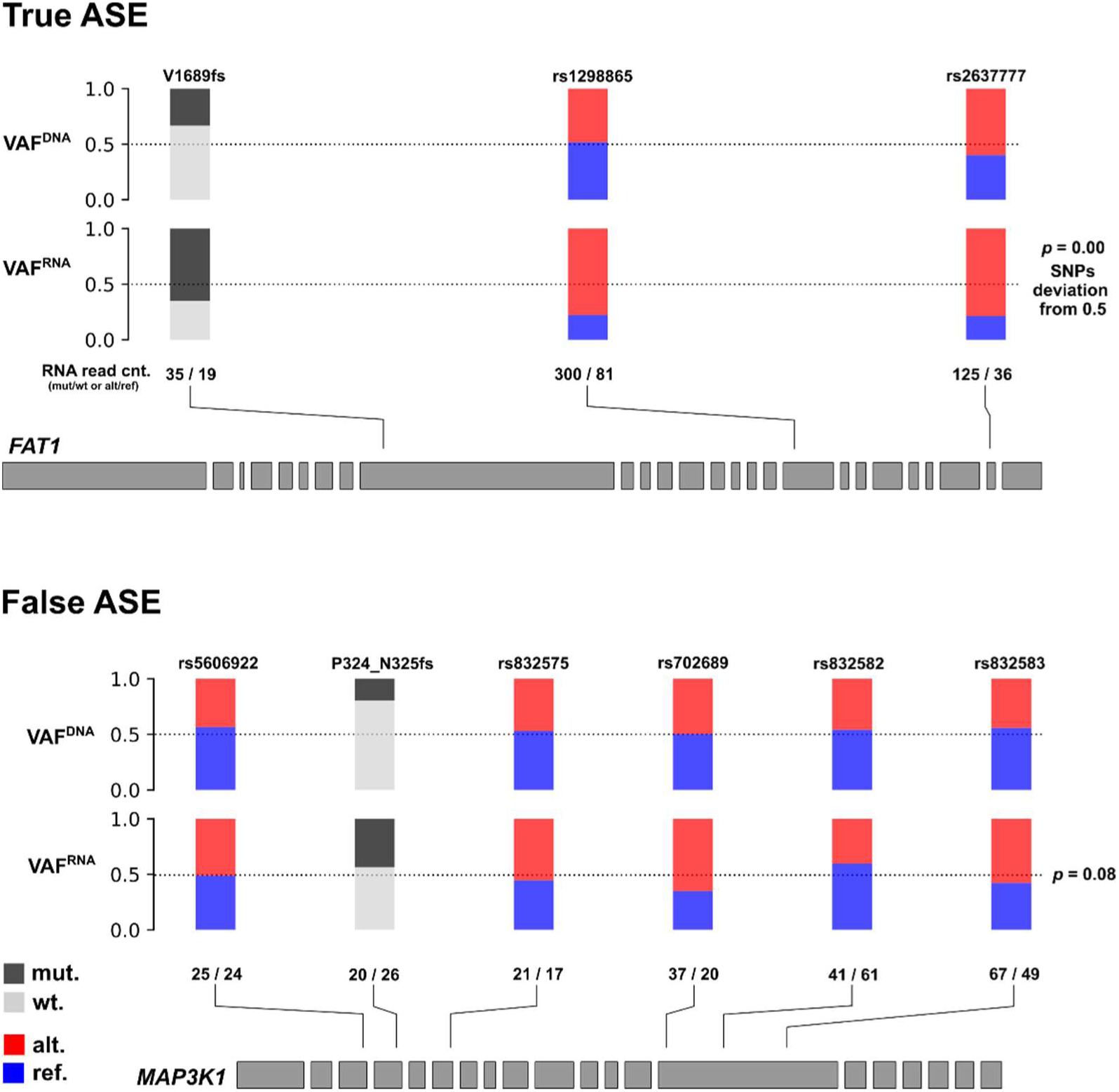
Distinguishing AEV caused by true ASE from that by confounding factors. The examples used here represent genes in diploid regions. *Top:* true ASE event with *FAT1* V1689fs in a head-and-neck squamous cell carcinoma (TCGA-DQ-5629) characterized by the SNP VAF^RNA^ deviating from 0.5 (excluding mutation VAF^RNA^) with a simulation-based *p* = 0.00 (Materials and Methods). *Bottom*: false ASE with *MAP3K1* P324_N325fs in a breast cancer (TCGA-BH-A0DT) exhibited an insignificant deviation with *p* = 0.08.

**Figure S6.**
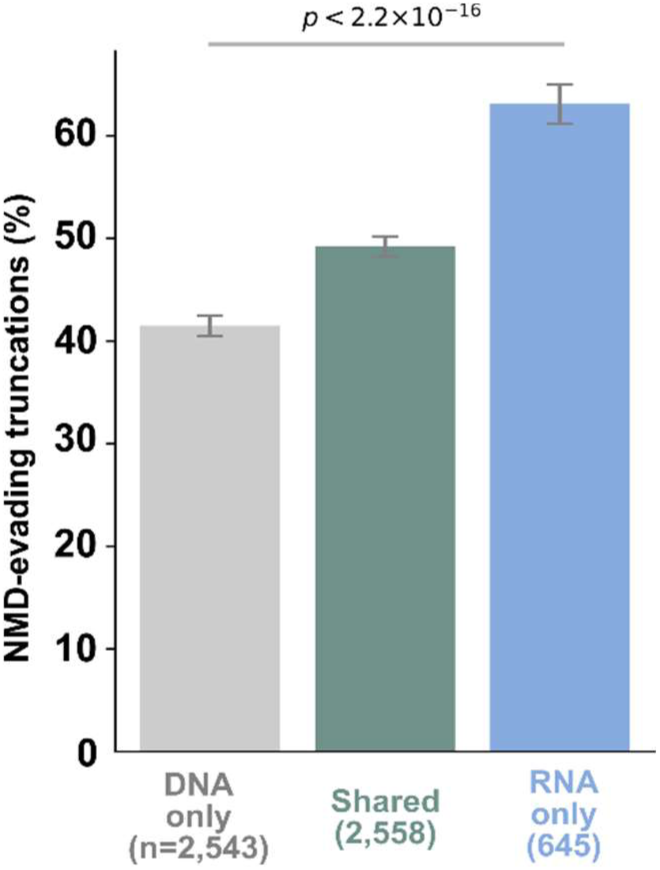
Frequency of NMD-evading truncating indels. The percentages of truncations in tumor suppressor genes with NMD evading features are plotted for each category. Comparison was made by χ^2^ test.

**Table S1.** Allele quantification benchmarking (DNA). Median VAF ratio and median absolute error (in parentheses) are shown at each dilution.

| Dilution | indelPost |  | MuTect2 |  | VarScan2 |  | Pindel |  |
| --- | --- | --- | --- | --- | --- | --- | --- | --- |
| 3/4 | 0.754 | (0.118) | 0.753 | (0.124) | 0.752 | (0.140) | 0.787 | (0.136) |
| 1/2 | 0.513 | (0.106) | 0.513 | (0.108) | 0.490 | (0.140) | 0.546 | (0.128) |
| 1/5 | 0.197 | (0.071) | 0.182 | (0.826) | 0.0 | (0.200) | 0.210 | (0.128) |
| 1/10 | 0.102 | (0.048) | 0.065 | (0.100) | 0.0 | (0.100) | 0.060 | (0.100) |
| 1/20 | 0.051 | (0.044) | 0.0 | (0.050) | 0.0 | (0.050) | 0.0 | (0.050) |

**Table S2.** Allele quantification benchmarking (RNA). Median VAF ratio and median absolute error (in parentheses) are shown for the ground truth genotypes. Het.: heterozygous (true VAF = 0.5). Hom.: homozygous (true VAF = 1.0)

| Genotype | indelPost |  | MuTect2 |  | VarScan2 |  | Pindel |  |
| --- | --- | --- | --- | --- | --- | --- | --- | --- |
| Het. | 0.450 | (0.125) | 0.400 | (0.183) | 0.037 | (0.474) | 0.000 | (0.5) |
| Hom. | 1.0 | (0.0) | 1.0 | (0.0) | 0.094 | (0.906) | 0.526 | (0.474) |

**Table S3.** Cost performance improvement by RNAIndel version 3. From renal clear cell carcinoma (TCGA KIRC), 10 RNA-Seq samples were selected ranging from 3.7 – 13.1 gigabyte (GB) data size. Refactored RNAIndel (version 3.0.1) [10] was compared to the original version (2.2.3) on Cancer Genomics Cloud [11]. The compute cost is based on the pricing as of Apr. 2021. USD: United States dollar.

| Case ID | BAM file size | version 2.2.3 |  | version 3.0.1 |  |
| --- | --- | --- | --- | --- | --- |
|  |  | Run time | Cost | Run time | Cost |
| TCGA-CZ-5465 | 12.3 GB | 356 min | 1.74 USD | 29 | 0.25 |
| TCGA-A4A57E | 5.2 | 86 | 0.53 | 11 | 0.09 |
| TCGA-CZ-5454 | 11.5 | 439 | 2.11 | 36 | 0.31 |
| TCGA-B9-4115 | 8.6 | 212 | 1.04 | 20 | 0.17 |
| TCGA-BQ-7059 | 10.3 | 365 | 1.79 | 40 | 0.34 |
| TCGA-BQ-7055 | 6.2 | 79 | 0.39 | 9 | 0.08 |
| TCGA-BQ-7045 | 3.7 | 42 | 0.26 | 7 | 0.06 |
| TCGA-CZ-5455 | 13.1 | 253 | 1.57 | 36 | 0.25 |
| TCGA-B0-5699 | 7.7 | 205 | 0.9 | 22 | 0.14 |
| TCGA-B0-4700 | 8.8 | 134 | 0.83 | 15 | 0.11 |

**Supplementary Tables S4–S6**

Tables S4–S6 can be found in Zenodo (https://zenodo.org/records/15492145).

